# Self-organized division of labor and selection for long-term antiviral defense in CRISPR arrays

**DOI:** 10.64898/2026.09.22.753571

**Authors:** Jimena Martín-Reina, Jaime Iranzo

## Abstract

CRISPR-Cas is an adaptive immunity system that protects prokaryotes from viruses and other genetic parasites. CRISPR-Cas immunity is guided by short DNA sequences, called spacers, that are acquired upon encounters with parasites and stored in the CRISPR array. Although the mechanisms driving spacer acquisition are relatively well understood, the timespan of CRISPR immune memory and, consequently, the very nature of CRISPR-Cas as a short-term or long-term defense mechanism remain controversial. Because CRISPR arrays have limited size, CRISPR-Cas systems face a fundamental trade-off between acquiring new spacers against recent infections and retaining old spacers for long-term defense. To investigate how CRISPR-Cas systems resolve this “fast response-versus-memory” trade-off, we developed a stochastic model that captures CRISPR array dynamics under varying abundances of endemic and episodic viruses. Simulations reveal three distinct evolutionary regimes that result in short-term, long-term, and “dual memory” arrays. These regimes are governed by the ratio of two measurable parameters: the length of the CRISPR array and the time required for transient epidemics to decay below basal endemic viral abundance. Spontaneous division of labor emerges in the dual memory regime, with spacers near the leader and distal ends specialized in short-term and long-term memory, respectively. Within-array division of labor results in a U-shaped longitudinal profile for the probability of finding the targets of CRISPR immunity among the local virome. Such profiles, that have been empirically observed in CRISPR arrays from the human gut microbiome, could inform future research on the environmental persistence of poorly characterized viromes.

## Introduction

Bacteria and archaea have evolved a wealth of defense systems to prevent infection by viruses, plasmids, and other mobile genetic elements [1, 2]. Among prokaryotic defense systems, CRISPR-Cas stands out for its adaptive nature, which provides its carriers with versatile and quickly evolvable immune repertoires [3, 4]. Complete CRISPR-Cas systems encompass two modules: the adaptation module and the effector module. Upon encountering genetic parasites, the adaptation module (consisting of *Cas1* and *Cas2*) can incorporate small fragments of the parasite’s DNA into the host’s CRISPR array. These fragments, called spacers, are transcribed and processed, becoming probes that allow identifying the parasite in future encounters. If identical (but reverse-complementary) sequences to those in the spacers are found, the effector module, consisting of diverse proteins depending on the CRISPR-Cas type, cleaves or silences them. The adaptation process occurs in a polar manner, with new spacers added to the leading end of the CRISPR array (the 5’ end in the direction of transcription) and separated from older spacers by short conserved sequences called repeats [5]. Random spacer loss can occur at any position of the array due to recombination among the conserved repeats that flank every spacer [6, 7]. Because of polar incorporation and random loss, CRISPR arrays provide an ordered, though incomplete, record of past encounters with genetic parasites.

Analysis of CRISPR arrays has proved a powerful strategy to decipher virus-host interactions, facilitating the discovery and characterization of novel viruses and other mobile genetic elements, especially from poorly studied environments [8-16]. From an ecological perspective, longitudinal tracking of CRISPR-mediated virus-host interactions is informative of microbial community dynamics [17, 18]. Environmental CRISPR surveys have revealed diverse CRISPR-Cas targeting profiles and activity levels among biomes, suggesting qualitative differences in the composition and persistence of the virome and the mobilome (that is, the population of mobile genetic elements in the community) [19-25].

Conceptual debates about the costs and benefits of CRISPR-Cas systems have traditionally revolved around their adaptive nature and its implications in terms of immune plasticity, specificity, and overall cost compared to other defense mechanisms [26, 27]. However, long-term immune memory (a property shared with the adaptive immune system of vertebrates) is also likely to play a pivotal role in the protective function of CRISPR-Cas systems. Long-term (months to years) maintenance of active spacers against locally persistent viruses and plasmids has been observed, for example, in microbial communities from acid mine drainages [28, 29], hypersaline lakes [19], and wastewater treatment plants [23]. In contrast, recent experiments with the model organism *Pseudomonas aeruginosa* PA14 (which carries a type I-F CRISPR-Cas system) and phage DMS3vir failed to induce long-lasting CRISPR-Cas immunity despite repeated exposure to the virus [30]. In fact, although long-term CRISPR memory is often observed in natural communities, it appears to be counter-selected in laboratory experiments. The ecological, evolutionary, and genetic factors that promote maintenance of long-term memory spacers in CRISPR arrays are still a matter of investigation, and likely include mobilome properties, such as virus and plasmid abundance, diversity, and endemicity, as well as host population-level properties, such as the strength of intraspecific and interspecific competition and the availability of other defense systems.

To be effective, CRISPR arrays must reach a balance between acquisition of new spacers and maintenance of immune memory [7, 31]. Spacer acquisition is necessary for coping with new parasites and counteracting escape mutations in the targets of CRISPR immunity. However, because arrays cannot grow indefinitely, faster spacer turnover comes at a cost of shorter memory spans. The large variability in the environmental persistence of different mobile genetic elements, from transient lytic viruses to endemic plasmids and prophages, raises the question of how CRISPR-Cas systems resolve the spacer turnover-vs-memory trade-off. It has been hypothesized that different arrays within a host could be specialized in long-term and short-term memory, which would explain the maintenance of multiple CRISPR arrays in a sizeable fraction of prokaryotic genomes [32, 33]. Alternatively, spacer-wise, age-dependent selection against loss of functional spacers could lead to self-organized division of labor within CRISPR arrays, with short-term memory spacers located in the leading region of the array and long-term memory spacers located in the distal region. Such division of labor has been observed in CRISPR arrays from the adult human gut microbiome [25], which is dominated by lysogenic phages and locally persistent crass-like phages [34-37].

In this article, we present a simple stochastic model that simulates the evolutionary dynamics of CRISPR arrays. By simulating CRISPR array dynamics under a broad range of conditions, we investigated the emergence of long-term CRISPR-Cas memory and identified which properties of the mobilome lead to fully specialized arrays (either for short-term or long-term memory) and which ones promote dual memory arrays. Conversely, we aimed at finding out what general properties of the local mobilome can be inferred by inspecting trends in target composition within CRISPR arrays. For simplicity, the presentation focuses on the episodic and endemic components of the local virome. Nevertheless, all the results can be extended to other components of the mobilome (including plasmids and integrative elements) by considering their transient or persistent nature in the host population.

## Model

### Virome properties and virus dynamics

We propose a simple model to study how the properties of the local virome determine the evolution and long-term composition of CRISPR arrays. The model considers two classes of viruses based on their infection dynamics at the population level: episodic and endemic. We assume that episodic (or transient) viruses appear once in the population, quickly reaching their peak abundance and then decaying exponentially at a rate *β* > 0 (the inverse of this rate, 1/*β* represents the characteristic time that the virus remains in the system). In contrast, endemic viruses persist for long periods of time with minimum decay (*β* = 0). To facilitate analytic calculations and interpretation of the results, we defined a derived parameter *b* = *e*^−*β*^ that represents virus persistence. This parameter takes values between 0 and 1, with *b* = 1 for pure endemic viruses and *b* ≪ 1 for episodic viruses that quickly decay in abundance. Given the absence of empirical measurements, we assumed that the persistence parameter of episodic viruses follows a uniform distribution between 0 and 1.

The properties of the local virome are determined by three additional parameters: *γ, ϕ*, and *α* (see Fig. 1A for a visual description).

**Figure 1.**
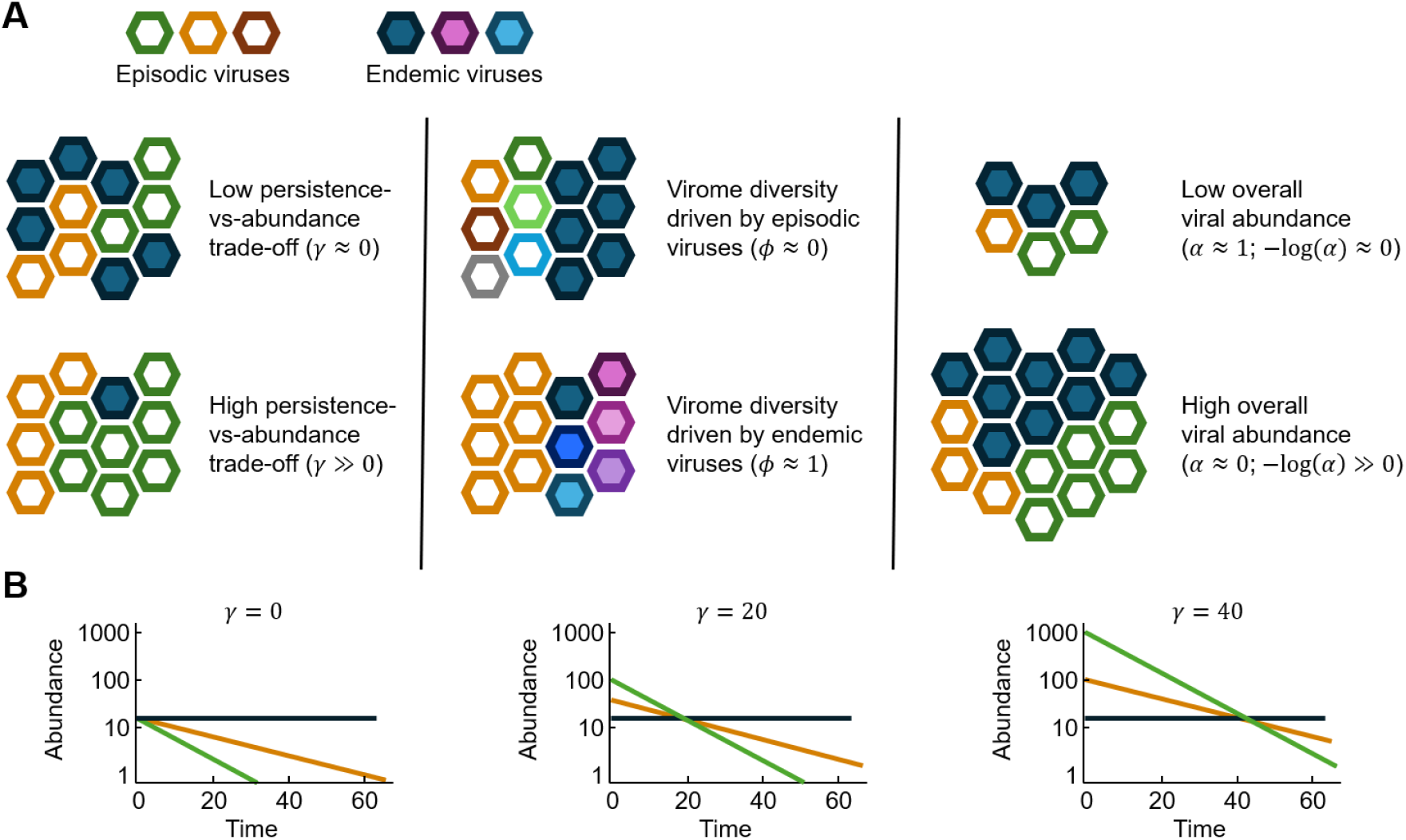
Key virome parameters determining the evolution of CRISPR arrays. A: Schematic representation of the persistence-abundance trade-off (parameter *γ*, left); the relative contribution of endemic viruses to the total virome diversity (parameter *ϕ*, center); and the overall virus abundance (parameter *α*, right). B: The persistence-abundance trade-off determines the time that it takes for episodic viral infections to decay from their peak abundances to the background abundance of endemic viruses. Therefore, parameter *γ* can also be interpreted as the characteristic duration of transient epidemics. Green and yellow lines correspond to two episodic viruses with different persistence indices (*b*_*yellow*_ > *b*_*green*_). Note the logarithmic scale for the viral abundances.

- Parameter *γ* controls the relative abundance of episodic viruses with respect to endemic viruses at the time of appearance. The model accounts for a possible trade-off between persistence and peak abundance, meaning that episodic viruses typically reach high peak abundances followed by fast decay, whereas endemic viruses persist at lower average abundances but for longer periods of time. Following this rationale, the peak abundance of a virus was modeled to be proportional to *e*^*βγ*^ (or, equivalently, *b*^−*γ*^), where *γ* = 0 represents the absence of a trade-off and *γ* ≫0 indicates much higher peak abundance of episodic viruses compared to endemic viruses. Note that, whereas *β* and *b* generally take different values in different viruses (see below), we used a single persistence-abundance trade-off *γ* to characterize the virome.
- Parameter *ϕ* defines the probability that new spacers target endemic viruses and is therefore directly related to the persistence of the virome. The extreme cases represent scenarios in which all new spacers correspond to episodic or endemic viruses (*ϕ* = 0 and *ϕ* = 1, respectively). Parameter *ϕ* also captures the effect of primed adaptation, since frequent reacquisition of spacers against endemic viruses results in higher values of *ϕ*.
- Parameter *α* controls the overall viral abundance or, equivalently, the selection pressure imposed by viruses on host survival. We parameterized *α* to be analogous to a selection index: *α* = 1 represents a neutral scenario in which viral infections are rare and viruses play a negligible role in bacterial population dynamics (in consequence, antiviral defense does not provide any selective advantage); in turn, *α* ≪ 1 represents a scenario in which viruses are abundant and infections are frequent, imposing a strong purifying selection to keep functional spacers in the CRISPR array. With this interpretation in mind, we defined the baseline viral abundance as log(1/*α*), with 0 < *α* ≤ 1.

Theoretical and empirical evidence suggests that spacer acquisition in natural environments is associated with peaks in viral abundance [25, 38]. Therefore, for the purpose of modeling CRISPR spacer dynamics, we assumed that the age of a spacer is equal to the time since its cognate virus reached its peak abundance. With the parameters defined above, the abundance of a virus at a time *t* after reaching the peak can be expressed as:

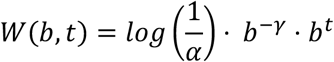

By rewriting the last two factors as *b*^*t*−*γ*^, it becomes evident that parameter *γ* admits an alternative interpretation as the typical time at which the abundances of endemic and episodic viruses become equal. Thus, after reaching their peak, episodic viruses prevail above the background of endemic viruses for a time window *t*^∗^ = *γ* (Fig. 1B).

### Spacer dynamics

The model tracks a lineage of CRISPR arrays subject to spacer gain, loss, and selection for host survival. For simplicity, we assumed that spacer gain and loss rates are at equilibrium, so that the number of spacers in the array, *N*, remains constant. For computational efficiency, selection was modeled as differential probability of loss for different spacers, under the assumption that the loss of spacers that target abundant viruses is deleterious and therefore such losses are less likely to reach fixation within the lineage.

We defined the dispensability *Q*_*i*_ of a spacer *i* as the probability that the host does not encounter the virus targeted by that spacer in a time unit. Assuming a Poisson process for virus-host encounters, the dispensability becomes 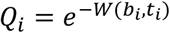, where *b*_*i*_ is the persistence of the virus targeted by spacer *i* and *t*_*i*_ is the age of that spacer.

With these considerations, simulations proceed as follows:

1. At each time step, a spacer *i* is added to the leading end of the array. Spacer acquisition represents the entry of a new virus in the system. With probability *ϕ*, the spacer is assigned to an endemic virus; with probability 1 − *ϕ*, it is assigned to an episodic virus. The persistence parameter of episodic viruses is randomly drawn from a uniform distribution in the interval (0,1), whereas the persistence parameter of endemic viruses is set to *b*_*i*_ = 0.9999 (we used that value instead of 1 for numerical stability).
2. Following spacer acquisition, we recalculate the dispensability of every spacer. Then a spacer *j* is randomly deleted with probability 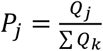, where *Q*_j_ is the dispensability of spacer *j*.

These steps were iterated until the spacer composition (that is, the distribution of *b*_*i*_ and *t*_*i*_ along the array) reached a stationary state.

As an extension of the basic model, we considered the possibility that mutations in viral protospacers render spacers ineffective in the long term. We assumed that the probability that a spacer matches its original protospacer exponentially decays with time as 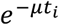, where μ is the protospacer mutation rate and *t*_*i*_ is the age of the spacer. Combining this expression with the expression for the target abundance *W*(*b*_*i*_, *t*_*i*_), the spacer dispensability becomes 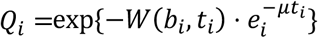.

A summary of the model parameters and their interpretation is provided in Table 1.

**Table 1.**
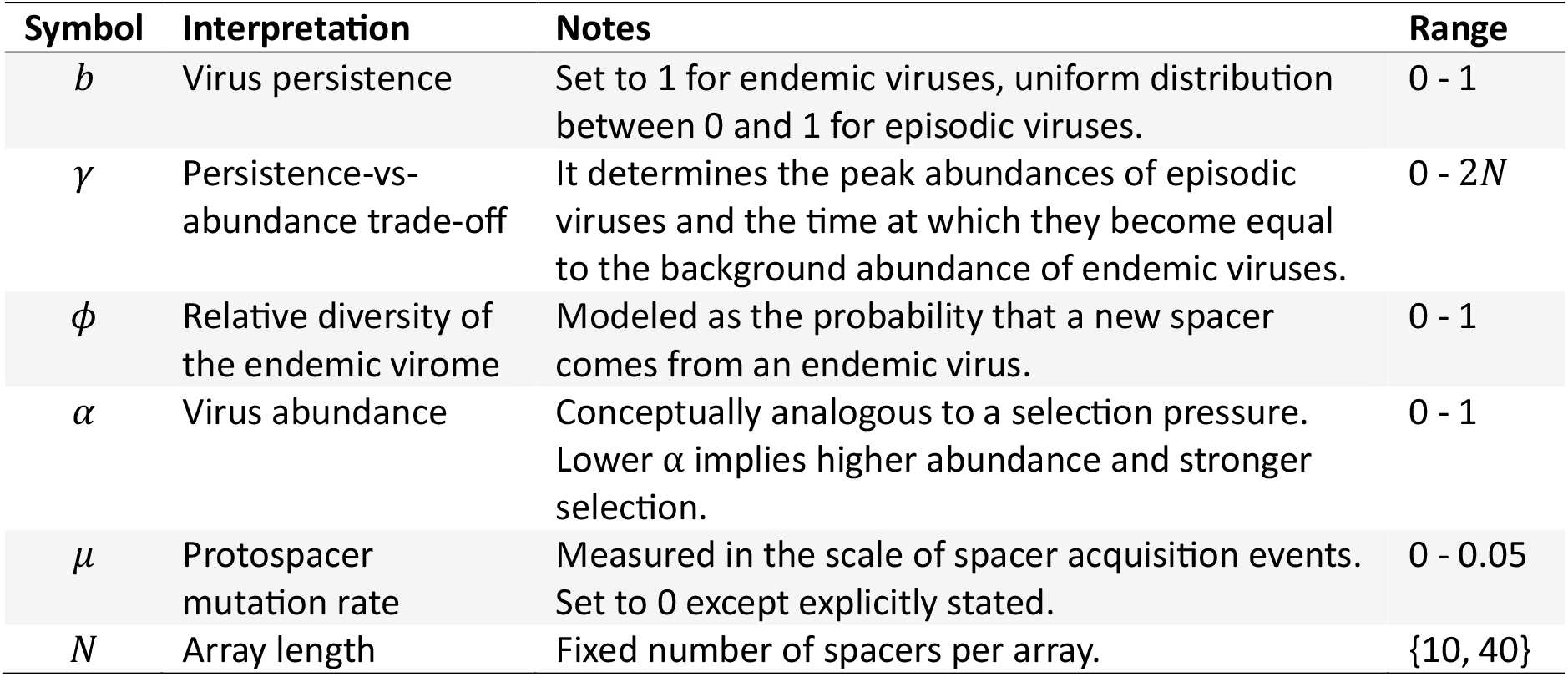
Model parameters.

| Symbol | Interpretation | Notes | Range |
| --- | --- | --- | --- |
| $b$ | Virus persistence | Set to 1 for endemic viruses, uniform distribution between 0 and 1 for episodic viruses. | 0 - 1 |
| $\gamma$ | Persistence-vs-abundance trade-off | It determines the peak abundances of episodic viruses and the time at which they become equal to the background abundance of endemic viruses. | 0 - $2N$ |
| $\phi$ | Relative diversity of the endemic virome | Modeled as the probability that a new spacer comes from an endemic virus. | 0 - 1 |
| $\alpha$ | Virus abundance | Conceptually analogous to a selection pressure. Lower $\alpha$ implies higher abundance and stronger selection. | 0 - 1 |
| $\mu$ | Protospacer mutation rate | Measured in the scale of spacer acquisition events. Set to 0 except explicitly stated. | 0 - 0.05 |
| $N$ | Array length | Fixed number of spacers per array. | {10, 40} |

## Results

### Distribution of long-term memory spacers within CRISPR arrays

To understand how the properties of the virome reflect on the composition of CRISPR arrays, we simulated the model under a broad range of parameter combinations and studied which kind of viruses (endemic or episodic) are targeted in each region of the array. In most cases, an asymmetric V-shaped trend emerges, with spacers that target persistent viruses decreasing in abundance from the leading end towards the middle of the array and then increasing again towards the distal end (Figs. 2 and S1). (The mean target persistence includes the contribution of episodic viruses, some of which could have relatively high values of *b*. Nevertheless, the curves for the fraction of spacers targeting strictly endemic viruses (*b* = 1) are very similar, except if *γ* > *N* and *ϕ* is small, in which case all spacers against endemic viruses are quickly lost.) The spacer at which target persistence profiles reach their minimum critically depends on the ratio between parameter *γ* (the strength of the persistence-vs-abundance trade-off) and the array length *N*. In fact, if *γ* = 0, that is, if the background abundance of endemic viruses is as high as the peak abundance of episodic viruses, the fraction of spacers that target endemic viruses monotonically increases from the leading end to the distal end of the array (Fig. 2, left panels). In contrast, if *γ* is greater than the length of the array (*γ* ≥ *N*, implying that the decay of episodic viruses is slower than the renewal rate of the full CRISPR array), the fraction of spacers that target endemic viruses decreases across almost the entire length of the array (Fig. 2, right panels). Finally, for intermediate values of *γ*, the fraction of the spacers that target endemic viruses follows the V-shaped curve presented above, with the minimum located near the *γ*-th spacer (Fig. 2, central panels). A systematic analysis of these curves confirms that the relative position of the minimum is approximately bounded by 1 − *e*^−*γ*/*N*^ and *γ*/*N* (Fig. 3), with values closer to the lower bound the greater the value of parameter *ϕ* (that is, the greater the diversity of endemic viruses compared to episodic viruses).

**Figure 2.**
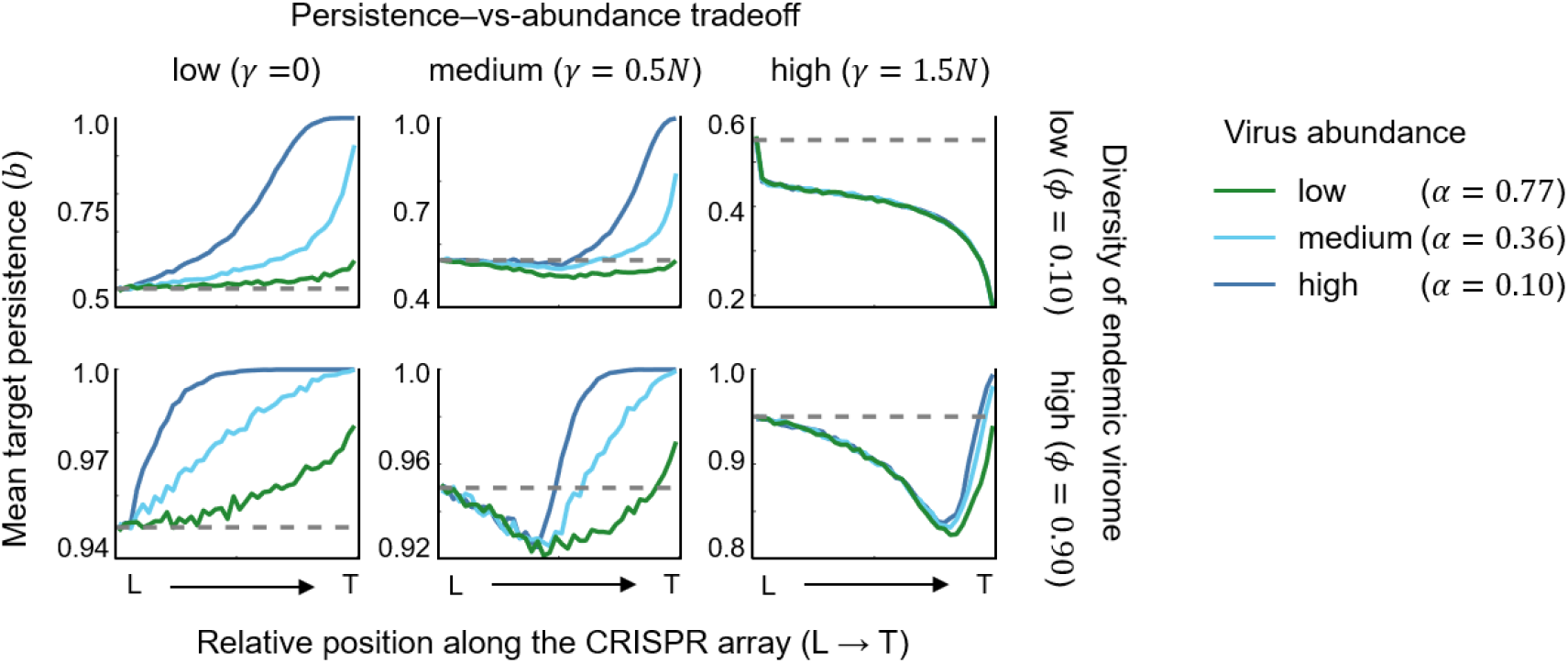
Spatial distribution of long-term memory spacers in the CRISPR array. Target persistence indices (1 for endemic viruses, less than 1 for episodic viruses, with lower values indicating faster decay) were recorded in each position of the array at the end of the simulation. The curves show averages across 10,000 simulations. We refer to these curves throughout the article as “target persistence profiles”. L: leader end of the array; T: trailing (distal) end of the array. The length of the array was set to *N* = 40 in all simulations. Dashed lines indicate the expected persistence in the absence of selection, which coincides with the average persistence of viruses targeted by the first spacer, (1 + *ϕ*)/2. See Fig. S1 for a broader range of parameter combinations.

**Figure 3.**
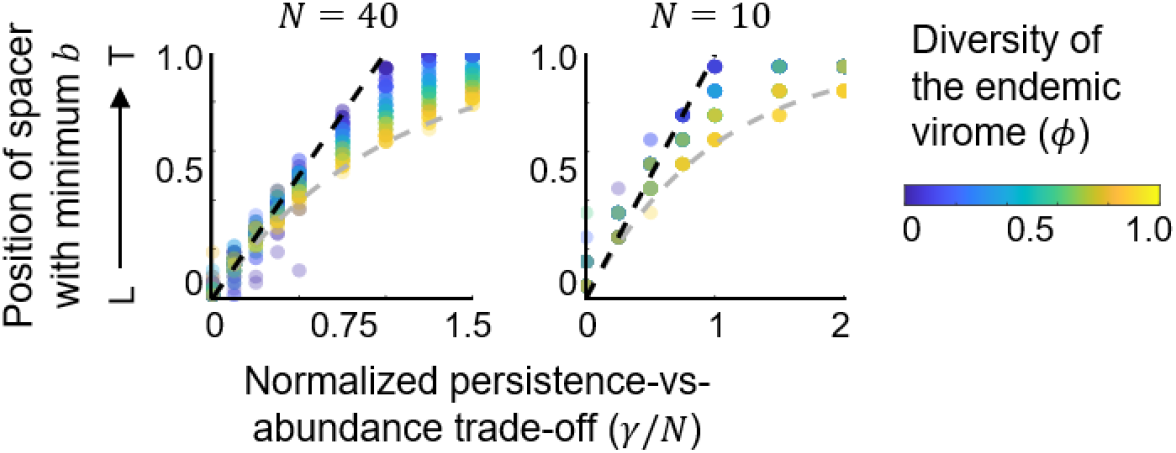
The persistence-vs-abundance trade-off defines short-term and long-term memory regions in the CRISPR array. The position of the array in which the target persistence profile reaches its minimum indicates the point where selection for long-term memory becomes dominant. These positions are plotted as a function of the persistence-vs-abundance trade-off *γ* (scaled by the array length *N*, x-axis) and the diversity of the endemic virome *ϕ* (color scale). The dashed lines represent two approximate boundaries for the observed points: *y* = *γ*/*N* (upper bound, black) and *y* = 1 − *e*^−*γ*/*N*^ (lower bound, gray). L: leader end of the array; T: trailing (distal) end of the array.

The V-shaped profile for the persistence of CRISPR targets results from two opposite selection pressures that operate at different time scales. In the short term, provided that the peak abundance of episodic viruses is greater than the background abundance of endemic viruses (*γ* > 0), selection will favor maintenance of spacers against episodic viruses. In the long term, once episodic infections have decayed, selection will favor the loss of such spacers. The time scale of the transition, *t*^∗^, is given by the abundance-vs-persistence trade-off. In fact, with the model parameterization, that time is simply *t*^∗^ = *γ* (Fig. 1B). By measuring time in terms of spacer replacement events, it becomes evident that the position of a spacer within the array determines a lower bound for its age (to move from the leader end to an arbitrary position *K*, at least *K* time steps are required). Therefore, all spacers after the *γ*-th position are older than *t*^∗^ and are subject to the long-term selection regime, which purges less useful spacers against episodic viruses. In contrast, selection in younger regions of the array favors immunity against episodic viruses, allowing for the progressive loss of spacers that target endemic viruses.

The conceptual link between overall viral abundance and selection for optimal spacer composition becomes clear when looking at the curves in Figs. 2 and S1. High viral abundances result in stronger selection and more efficient accumulation of spacers against endemic viruses towards the distal end of the array. If selection is strong enough, spacers against episodic viruses are completely purged, generating a distal region that is fully adapted for long-term memory (right plateaus for *α* = 0.1 in Fig. 2). Such fully specialized regions do not appear if selection is weak, although a partial enrichment in spacers against persistent viruses is still observed in those cases, provided that the decay of episodic viruses is not too slow (that is, provided that *γ* < *N*, Fig. S2). Somewhat paradoxically, selection for memory spacers against environmentally persistent viruses does not necessarily imply that distal spacers are older than expected under a neutral scenario (Fig. S3). In fact, long-term retention of memory spacers is only observed if *γ* is substantially smaller than *N*, that is, if most of the array is specialized in defense against endemic viruses.

### Contribution of spacers to local adaptation

The contribution of a spacer to CRISPR-Cas immunity depends on how likely it is for the host to encounter the spacer’s target in the local population. That, in turn, depends on the dynamics of the target (episodic or endemic) and, in the case of episodic infections, the time since the spacer was acquired. Because both the nature of the targets and the age of the spacers vary along the CRISPR array, different regions of the array are likely to contribute to varying degrees to the overall CRISPR-Cas immunity. To assess these differences, we obtained, for each spacer in the array, the probability that the host encounters the targeted virus over one timestep (Figs. 4 and S4). As expected, the probability of encountering the target strongly depends on the overall viral abundance, with higher viral loads (lower *α*) resulting in more frequent targeting. For most parameter combinations, the probability to encounter the target displays a minimum at an internal spacer and plateau-like maxima at both ends. Qualitatively, this implies that the most useful spacers (those with the greatest contribution to immunity) are located on both ends of the CRISPR array, whereas central spacers play a lesser role in immunity. In agreement with the target persistence profiles, the position of the minimum is proportional to parameter *γ* and its prominence depends on parameter *ϕ*, with deeper minima found in viromes with less diverse endemic components (lower *ϕ*). The plateau-valley-plateau trend results, on one side, from the decline in abundance of episodic viruses, which leads to a quick drop in the probability of finding the target as the age of spacers reaches the critical time *t*^∗^ = *γ*, and, on the other side, from delayed selection for spacers against endemic viruses, which drives the recovery of immune efficacy in distal regions of the array, where such spacers become dominant. In consequence, the minimum value is determined by the prevalence of spacers against endemic viruses at the critical time *t*^∗^ = *γ*, before selection for these spacers comes into play (hence its dependence on *ϕ*). The width of the valley is determined by the strength of selection: the stronger, the faster the enrichment in spacers against endemic viruses and the concomitant recovery of immune efficacy.

**Figure 4.**
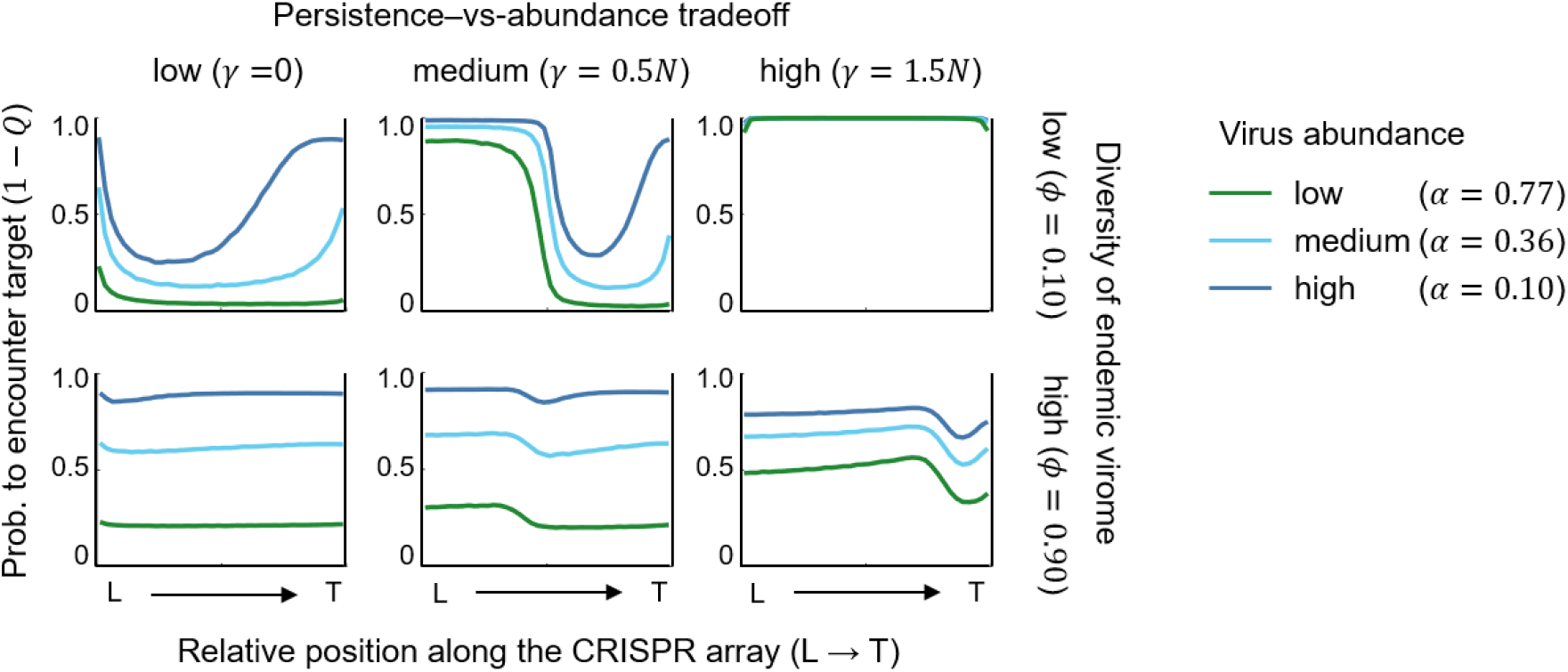
Contribution of different regions of the array to CRISPR-Cas immunity. The probability of encountering the target in one time step (given by the expression 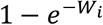, where *W*_*i*_ is the target abundance or, equivalently, 1 − *Q*_*i*_, where *Q*_*i*_ is the spacer dispensability) was recorded for every spacer at the end of the simulation. The curves show the medians of 10,000 simulations. We refer to these curves throughout the article as “target detection profiles”. L: leader end of the array; T: trailing (distal) end of the array. The length of the array was set to *N* = 40 in all simulations. See Fig. S4 for a broader range of parameter combinations.

### Selection for long-term memory is weakly affected by mutations in protospacers

The results presented so far were obtained under the assumption that spacers maintain their functionality over time. However, mutations in protospacers could let viruses escape recognition by CRISPR-Cas effector complexes, turning old spacers ineffective (and therefore, dispensable). In some CRISPR-Cas subtypes, partially matching spacers can still contribute to immunity by facilitating the acquisition of new (updated) spacers, through a mechanism known as primed adaptation. But even in those cases, partially matching spacers would become dispensable once the updated spacers have been acquired. In this section, we investigate whether selection for long-term memory spacers can operate despite escape mutations.

The spontaneous mutation rates of DNA viruses are in the order of 10^-8^ to 10^-6^ substitutions per base per round of replication, with values in the lower end corresponding to small viruses, such as ΦX174, and values in the higher end to large bacteriophages, such as T4 [39-41]. When scaled to protospacers of 30-40 nucleotides, the substitution rate per target becomes of the order of 10^-7^ to 10^-5^ per round of replication, although higher values are possible in mutator strains. To include these values in the model, we would need to know how the rate of spacer acquisition (that determines the unit of time in the model) compares to viral replication rates. Estimating the number of virus replication rounds between two consecutive spacer acquisitions (not necessarily targeting the virus of interest) is not trivial. As a workaround, we tested a broad range of values reaching up to 0.05 mutations per protospacer per time step, which, given the mutation rates presented above, would cover from 1 up to 5,000-500,000 rounds of division per spacer acquisition, for small and large viruses, respectively.

The results of an expanded model that accounts for mutations in protospacers are shown in Fig. 5. In general, nonzero mutation rates have limited effect on the target persistence profiles. Major drops in the prevalence of long-term memory spacers become apparent only at the highest mutation rate (*μ* = 0.05). Even in those extreme cases, persistence profiles show a minimum at approximately the *γ*-th spacer, after which the fraction of spacers against endemic viruses increases. Overall, the profiles obtained for increasing mutation rates are qualitatively similar to those for decreasing viral abundances (compare curves in Fig. 5 with those for *γ* = 0.5 *N* and varying *α* in Fig. 2). Because of the conceptual connection between virus abundance and selection strength, the resemblance across profiles for varying *μ* and varying *α* is consistent with the interpretation that mutations in protospacers result in weaker selection for long-term memory spacers, though the magnitude of this effect is small even with moderate mutation rates.

**Figure 5.**
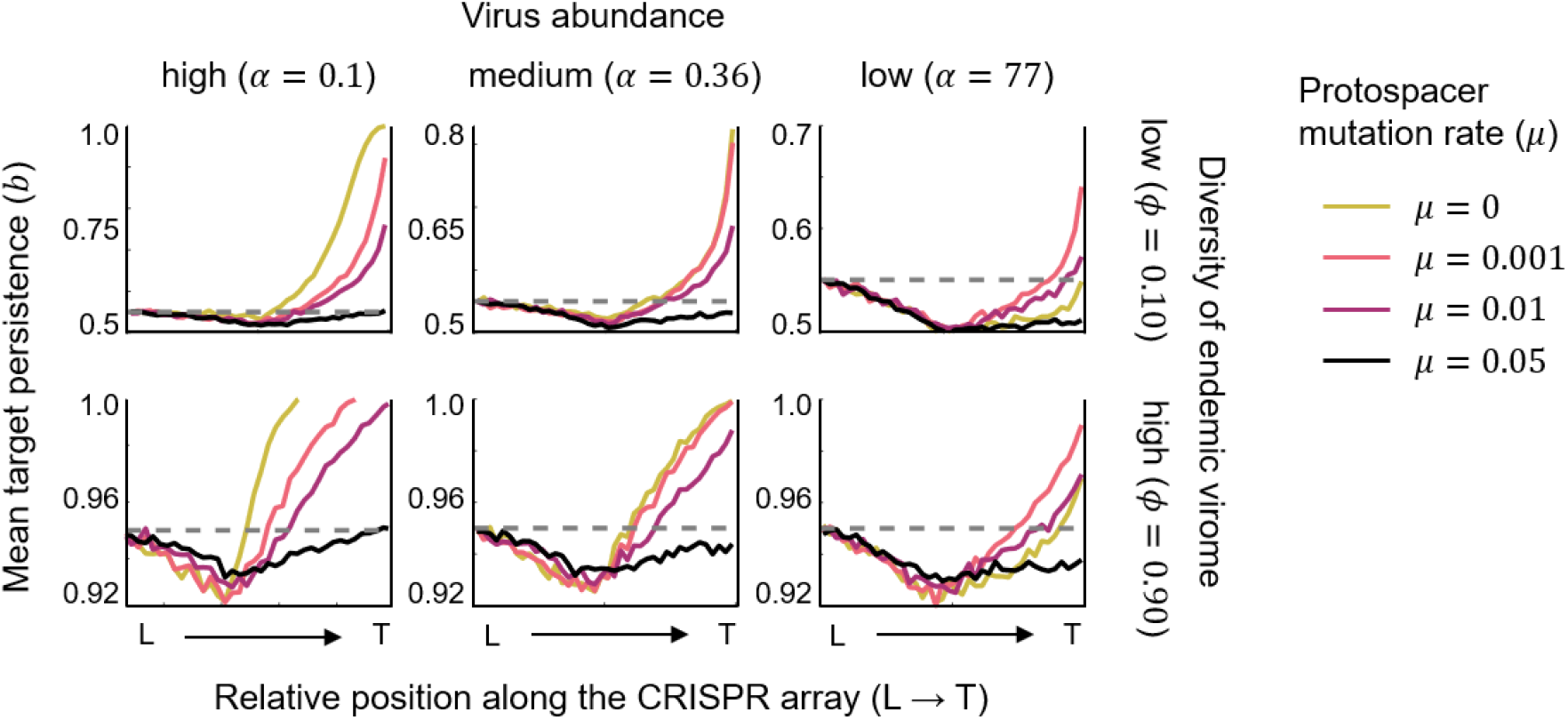
Effect of protospacer mutations on target persistence profiles. Target persistence indices (1 for endemic viruses, less than 1 for episodic viruses, with lower values indicating faster decay) were recorded in each position of the array at the end of the simulation. The curves represent averages across 10,000 simulations. Results are shown for 3 values of the virus abundance (left, center, and right panels) and 2 values of the endemic diversity (top and bottom panels). L: leader end of the array; T: trailing (distal) end of the array. The length of the array and the persistence-vs-abundance trade-off were set to *N* = 40 and *γ* = 20, respectively, in all simulations. Dashed lines indicate the expected persistence in the absence of selection, which coincides with the average persistence of viruses targeted by the first spacer, (1 + *ϕ*)/2.

## Discussion

We explored a simple model of CRISPR array dynamics to investigate how long-term immune memory emerges and is maintained within CRISPR arrays. In agreement with empirical observations, we identified three distinct evolutionary regimes leading to short-term, long-term, and “dual memory” allocation. Remarkably, these three regimes can be explained by the ratio of two measurable parameters: the time it takes for transient epidemics to decay below the basal abundance of endemic viruses (represented in the model by the persistence-vs-abundance trade-off) and the length of the CRISPR array.

For sufficiently long arrays and moderate trade-offs, selection for antiviral efficacy leads to a spontaneous division of labor within the CRISPR array. In these dual memory arrays, short-term defense against episodic viruses is carried out by recently acquired spacers in the leading region of the array, whereas long-term memory against persistent viruses concentrates in older, distal regions of the array. Division of labor is less prominent or not observed if episodic viruses remain abundant for long periods of time (longer than the characteristic renewal time of the whole CRISPR array). This condition, that can be fulfilled if the persistence-vs-abundance trade-off is sufficiently large or if arrays are short and spacer turnover is fast, leads to selection for short-term immunity across the whole array. On the opposite end, in environments in which the peak abundance of episodic viruses does not surpass the basal abundance of endemic viruses (no persistence-vs-abundance trade-off), selection leads to enrichment in long-term memory spacers along the whole CRISPR array.

Although the distribution of virus decay times is a property of the virome, the fact that timescales depend on spacer and array renewal rates gives hosts some control over memory allocation. Increasing or reducing the spacer turnover rate is effectively equivalent to extending or shortening persistence times, respectively. Accordingly, full array specialization in short-term memory can be achieved by reducing the size of the array and increasing the spacer acquisition rate. In turn, specialization in long-term memory can be achieved by reducing the spacer acquisition rate. As long as CRISPR arrays with different sizes and turnover rates can coexist in the same host [33], harboring long-term and short-term memory arrays appears as a feasible solution to deal with heterogeneity in virus persistence, as previously hypothesized [32]. Whether such distributed solution is preferable to within-array division of labor likely depends on multiple factors, including the relative cost of maintaining and expressing multiple arrays (possibly associated with different Cas operons) compared to a single longer array [42, 43], the degree of transcription of distal spacers [44], the intracellular availability of effector complexes [31], and the presence of alternative (non-adaptive) defense mechanisms [45, 46].

The immediate utility of a spacer is given by the probability of encountering its target. Therefore, the target detection profiles in Figs. 4 and S4 are informative of the intensity of selection against spacer loss in different regions of the array. The shape of these profiles, with wide plateaus and a central valley of variable depth and location, could explain the recent finding that effective loss rates are homogeneous along the CRISPR array [47]. In fact, if arrays from different environments and CRISPR-Cas types were pooled, as in the referred study, the resulting detection profile would become approximately flat except in both ends, where higher utility would result in lower effective loss rates.

A central prediction of the model is that selection can shape the content of CRISPR arrays by preventing the loss of useful spacers. Indeed, the strength of selection, which in the model is related to the overall viral abundance, is the strongest predictor of the enrichment in long-term memory spacers at the distal end of the array. Empirical evidence for selection on CRISPR array content includes the observation that CRISPR spacers preferentially target core viral and plasmid genes [13, 48], although these findings could be partly explained by preferential acquisition of spacers from early injected genomic regions, as shown for type II CRISPR-Cas and *Staphylococcus* phage Φ12 [49] and for type I-E CRISPR-Cas and MOB_F_ plasmids [50]. The role of selection in promoting long-term CRISPR memory is further supported by the recurrent observation that spacer diversity decays along the array [51] and distal spacers often target conserved regions of the genomes of persistent viruses [28, 52]. Based on our results, we propose that target detection profiles along CRISPR arrays can be used to assess selection for long-term CRISPR-Cas memory, with U-shaped profiles indicating strong selective maintenance of memory spacers. In environments in which information about target persistence is available (for example, through long-term virome surveys or detection of integrases as proxies for lysogenic lifestyle), ascendent or descendent trends in target persistence profiles provide additional information about selection for long-term or short-term memory, respectively. U-shaped target detection profiles combined with ascendent target persistence profiles have been recently reported by a large-scale study of CRISPR arrays in the adult human gut [25]. Such profiles suggest that persistent phages are dominant and impose a sustained selection pressure on the composition of CRISPR arrays, which fully agrees with the long-term stability and high abundance of crass-like phages in the adult human gut virome [34-37]. Considering the increasing availability of environmental metagenomes and bioinformatic tools for CRISPR detection and assembly [53], we believe that position-specific target profiling of CRISPR arrays can become a powerful yet simple strategy to shed light on the ecology and stability of the viromes of lesser studied environments.

### Limitations of the study and future prospects

The eco-evolutionary conditions required for the spread and maintenance of CRISPR-Cas in a host population have been the subject of much theoretical and experimental work (reviewed in [54-56]). To keep the model simple, we assumed that such conditions are held throughout the simulation. Theory and *in vitro* experiments suggest that CRISPR-Cas becomes inefficient if the diversity of the virome is too large [28, 57-59], a prediction that is consistent with the negative association between CRISPR-Cas prevalence and virus diversity observed in some biomes [60]. We investigated the effect of varying the relative contribution of episodic and endemic viruses to the overall virome diversity, while assuming that the absolute diversity is compatible with effective CRISPR-Cas defense. Provided that the absolute virome diversity determines CRISPR-Cas efficacy, we hypothesize that the effect of the absolute virome diversity on CRISPR memory can be captured by a change in the selection strength (parameter *α* in the model). Specifically, higher diversity would lead to lower overall CRISPR-Cas efficacy and, therefore, weaker selection on the CRISPR array content (*α* closer to 1).

For simplicity, we did not consider the possibility that expression levels decay along the array, which would reduce the efficacy of distal spacers. A decrease in transcription due to early termination has been reported in some CRISPR-Cas systems [61, 62]. In others, internal promoters and antitermination mechanisms facilitate transcription of the full array [63, 64]. Reduced transcription of distal spacers would have two quantitative effects on the array content profiles. First, early termination would result in an effective array length shorter than the actual length, *N*_*e*_ < *N*, where *N*_*e*_ is the position at which expression becomes negligible. The fitness contribution of every spacer after that position would be null, regardless of whether they targeted episodic or endemic viruses. As a result, target persistence profiles would level off after the *N*_*e*_ -th spacer, with no further enrichment in endemic or episodic targets. Second, because of their null contribution to fitness, selective loss of spacers would occur with highest probability in distal regions, resulting in relaxed purifying selection on the rest of the array. Therefore, before the *N*_*e*_ -th spacer, target persistence and spacer efficacy profiles would resemble those obtained with the current model under weaker selection (greater *α*).

Another major simplification of the model was to consider a fixed length for the CRISPR array. Array length distributions compiled from complete genomes and metagenomes are characterized by high variability and long tails [65], whereas those from populations evolved *in vitro* display a much narrower range of values [66, 67]. The long-tailed distributions observed in nature could be either intrinsic to the spacer gain and loss dynamics [65, 68] or the result from aggregating data from independent populations of arrays, each evolving under different acquisition and lost rates and distributed around a well-defined mean value [31, 44, 69]. In the latter case, small variations around the mean would be unlikely to change the main conclusions of the model. If size heterogeneity was, however, an intrinsic property of CRISPR array populations, we could speculate about a scenario in which division of labor did not only occur within single CRISPR arrays, but at the whole population level, with the shortest arrays specialized in short-term memory and longer arrays performing long-term or dual memory functions. In that scenario, population-level heterogeneity in CRISPR array lengths could represent a bet-hedging strategy to deal with the heterogeneous timescales of viral infections. More detailed models integrating population-level effects, eco-evolutionary feedback between CRISPR repertoire size and virome diversity, and empirical estimates of intra-population CRISPR diversity will be required to investigate this possibility.

## Methods

The source code (in C++ and Python) and datasets used in this study are available in GitHub [https://github.com/jimenamartinreina/modeling-crispr-arrays]. For each parameter combination, 10,000 random simulations were run to collect final summary statistics. To ensure that each simulation had reached the stationary state, we implemented a dynamic stop criterion. First, we ran the model for 5,000 time steps (*T*_*max*_) and checked the age of the oldest spacer. If the age of that spacer was >0.2 × *T*_*max*_, we doubled the maximum number of iterations and continued the simulation until the condition was no longer held. This strategy ensures that, by the time the simulation ends, any trace of the initial condition has been lost and the system has had time to settle into stationary state. For each parameter combination, we collected the following variables in each position of the array (*i*), averaging over replicates: the age of the spacer (*t*_*i*_), the persistence index of the target virus (*b*_*i*_), the target abundance (*W*_*i*_), the dispensability of the spacer (*Q*_*i*_), and the fraction of replicates in which the target is endemic.

For extreme parameter combinations (e.g. *γ* = 60 and *N* = 40), viral abundances can become very large, causing numerical overflow. To prevent this, abundance values were stored on a logarithmic scale and the dispensability was computed as 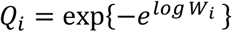 only if the abundance was below a threshold (log *W*_*i*_ < 40). Above that threshold, the dispensability was set to zero, since the exponential of very large negative values is functionally indistinguishable from zero in this context. We compared the results of the model with and without this implementation and the differences were indistinguishable from sampling noise.

## Acknowledgements

This work was funded by MICIU/AEI/10.13039/501100011033 and by ERDF/EU (PGE) “A way of making Europe”, through grants PID2019-106618GA-I00 and CNS2023-145430 to J.I. and by a JAE Intro fellowship (ref. JAEINT24_EX_1391) from the Spanish National Research Council (CSIC) awarded to J.M.R.

## Author contributions

J.I. designed the study; J.I. and J.M.R. developed and implemented the model; J.M.R. ran the simulations. All authors analyzed the results and wrote the manuscript.

