## Supplementary Figures for "Self-organized division of labor and selection for long-term antiviral defense in CRISPR arrays"

[1] Centro de Astrobiología (CAB), CSIC-INTA, Torrejón de Ardoz, Spain.
[2] Centre for Biocomputation and Physics of Complex Systems (BIFI), Zaragoza, Spain.

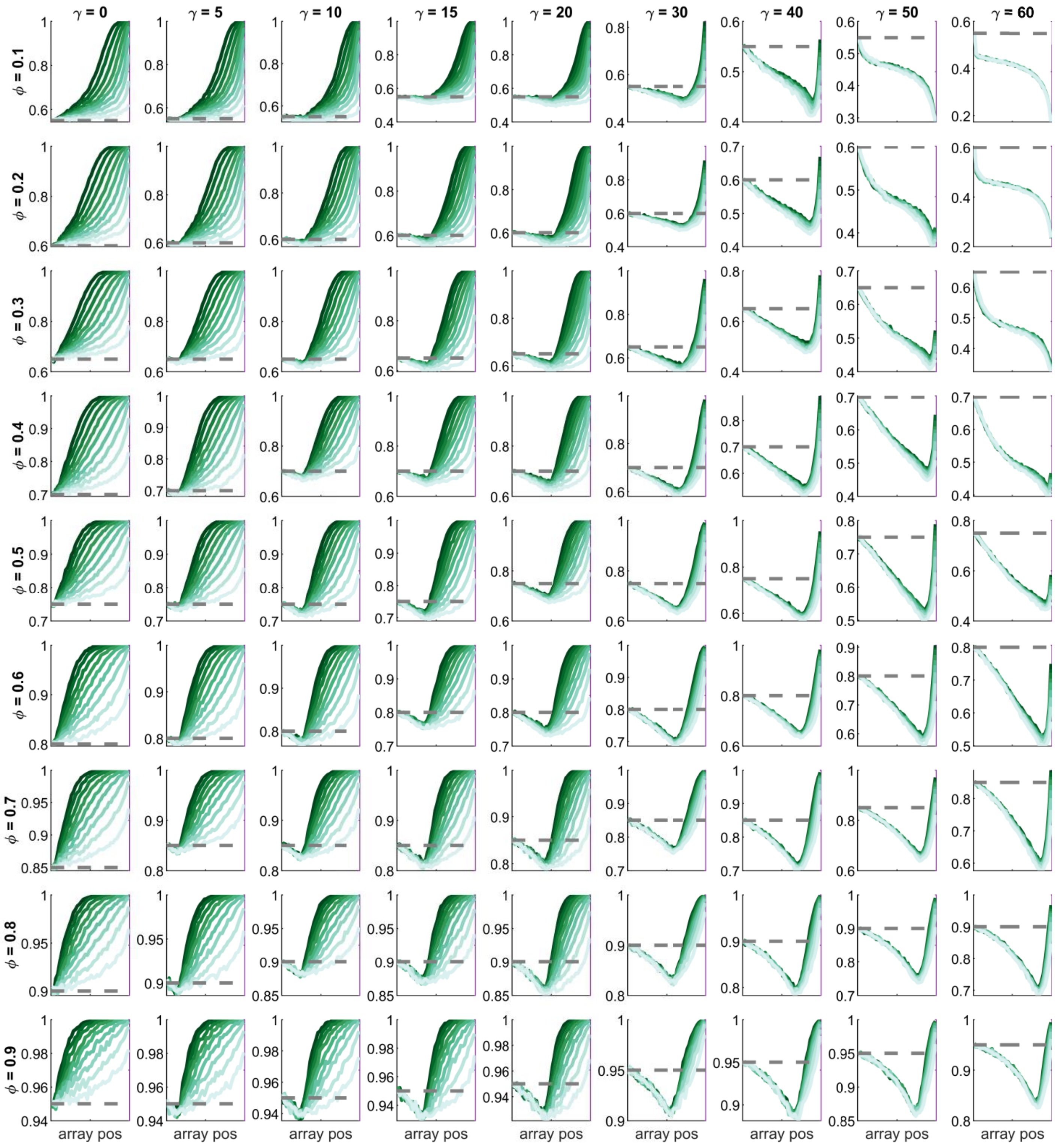


**Figure S1: Target persistence profiles along the CRISPR array.** Average target persistence index calculated over 10,000 simulations of the model (y-axis) as a function of the position of the spacer in the array (x-axis). Panels represent different combinations of the parameters $\phi$ (the relative contribution of endemic viruses to the overall virome diversity) and $\gamma$ (the strength of the abundance-vs- persistence trade-off). Curves of different colors represent different overall virus abundances, from least abundant (brightest color, minimum $\alpha = 0.77$) to most abundant (darkest color, $\alpha = 0.1$). The size of the array was fixed to $N = 40$ in all simulations. Dashed lines indicate the expected persistence in the absence of selection, which coincides with the average persistence of viruses targeted by the first spacer ($\frac{1+\phi}{2}$).

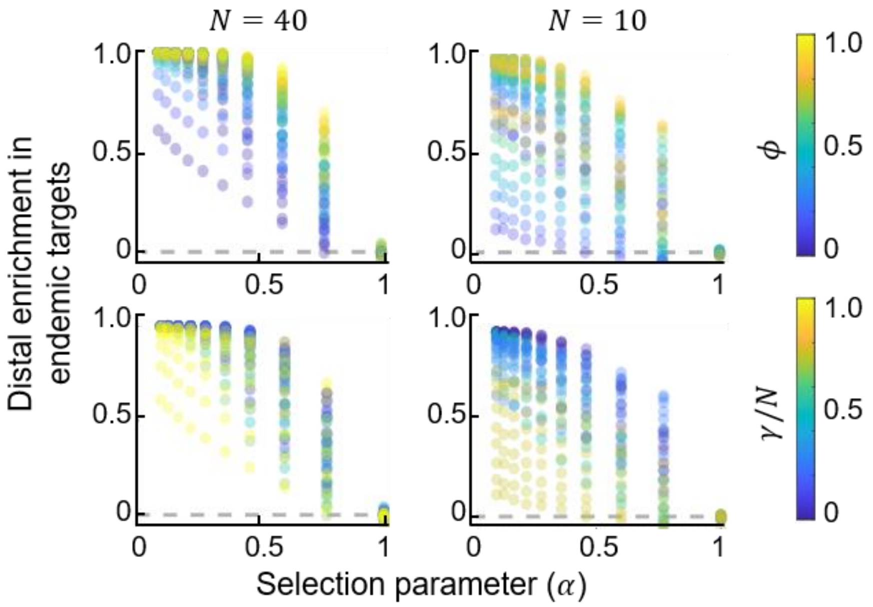


**Figure S2: Distal enrichment in endemic targets as a function of the model parameters.** The enrichment in endemic targets was calculated as $(\langle b \rangle - b_0)/(1 - b_0)$ where $\langle b \rangle$ is the target persistence index of the most distal spacer averaged over 10,000 simulations and $b_0 = (1 + \phi)/2$ is the expected value in the absence of selection. Each plot corresponds to a unique parameter combination. Gray dashed lines indicate no enrichment in endemic targets. Both rows show the same data, but with different coloring schemes.

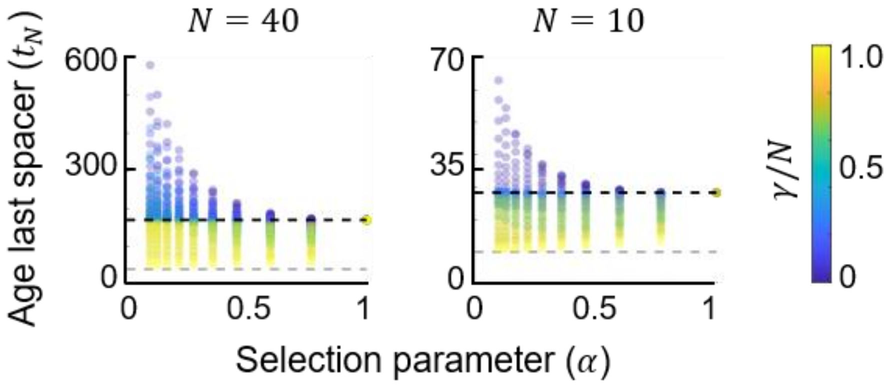


**Figure S3: Age of the most distal spacer in the array as a function of the model parameters.** Each point corresponds to a unique parameter combination. Gray dashed lines correspond to the minimum possible age for the last spacer, which is given by the length of the CRISPR array ($t_N = N$). Black dashed lines indicate the age of the most distal spacer in the absence of selection and were empirically obtained from simulations with $\alpha = 1$.

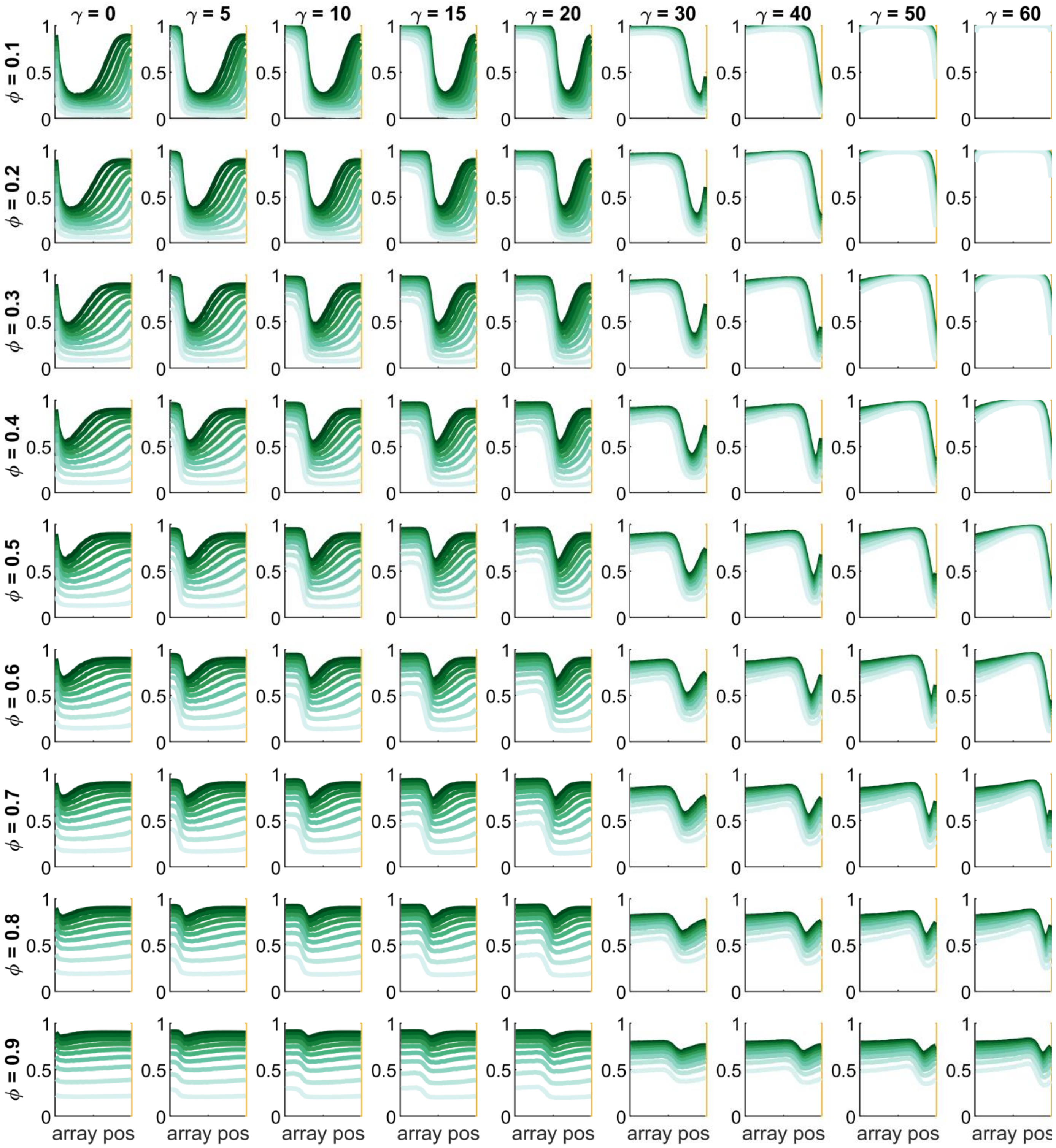


**Figure S4: Target detection profiles along the CRISPR array.** Median probability of encountering the target, calculated over 10,000 simulations of the model (y-axis) as a function of the position of the spacer in the array (x-axis). Panels represent different combinations of the parameters $\phi$ (the relative contribution of endemic viruses to the overall virome diversity) and $\gamma$ (the strength of the abundance-vs- persistence trade-off). Curves of different colors represent different overall virus abundances, from least abundant (brightest color, minimum $\alpha = 0.77$) to most abundant (darkest color, $\alpha = 0.1$). The size of the array was fixed to $N = 40$ in all simulations.
